# Droplets of amyotrophic lateral sclerosis-associated p62/SQSTM1 mutants show slower inner fluidity

**DOI:** 10.1101/2021.06.07.447422

**Authors:** Faruk Mohammad Omar, Yoshinobu Ichimura, Shun Kageyama, Afnan H. El-Gowily, Yu-shin Sou, Masato Koike, Nobuo N Noda, Masaaki Komatsu

## Abstract

A series of amyotrophic lateral sclerosis (ALS)-related proteins such as FUS, TDP-43 and hnRNPA1 has an ability to be liquid-liquid phase separation, and their disease-related mutations cause the transition of their responsible liquid droplets to aggregates. Missense mutations in *SQSTM1/p62*, which have been identified throughout the gene, are associated with ALS, frontotemporal degeneration (FTD) and Paget’s disease of bone. SQSTM1/p62 protein forms liquid-droplets through the interaction with ubiquitinated proteins, and the droplet serves as a platform of autophagosome formation and anti-oxidative stress response via the LC3-interacting region (LIR) and Keap1-interacting region (KIR), respectively. However, it remains unclear whether ALS/FTD-related p62 mutations in LIR and KIR form aberrant liquid droplets, cause defective autophagy and stress response or both. To evaluate the effects of ALS/FTD-related p62 mutations in LIR and KIR on a major oxidative stress system, the Keap1-Nrf2 pathway and the autophagic turnover, we developed systems that enable to monitor them with high sensitivity. These systems revealed that some mutants but not all have their less abilities on the Nrf2-activation and show the delayed turnover. By contrast, while the sufficient ability to form liquid droplets, all droplets consisting of p62 mutants showed slower inner fluidity. These results indicate that like other ALS-related mutant proteins, a primary defect in ALS/FTD with p62 missense mutations is a qualitative change of p62-liquid droplets.

## Introduction

*SQSTM1*, the human gene encoding for p62/SQSTM1 (hereafter as p62) localizes in chromosome 5 and comprises 8 exons distributed through 16 kb. Missense mutations in *SQSTM1* have been primarily associated with amyotrophic lateral sclerosis (ALS), frontotemporal lobar degeneration (FTD) and Paget’s disease of bone (PDB) (1,2). Different p62-positive structures such as inclusion bodies have been identified in patients suffering from neurodegenerative diseases including ALS, and FTD (3–5). However, a common mechanism how each mutation recognized throughout the gene encoding p62 protein causes ALS and FTD remains unclear.

p62, initially identified as a 62 kDa protein, has multiple functional domains which include an N-terminal Phox1 and Bem1p (PB1) domain, a zinc finger (ZZ), a tumor necrosis factor receptor-associated factor 6 (TRAF6) binding (TB) motif, an LC3-interacting region (LIR), a Keap1-interacting region (KIR) and a ubiquitin-associated (UBA) domain (6,7). The protein localization is not limited to the cytoplasm, but can also be observed in the nucleus, in autophagosomes and on lysosomes (8–10). Due to its role as an adaptor for selective autophagy, p62 also localizes to cargos that will be degraded during the process, such as ubiquitin-positive protein aggregates, damaged mitochondria and invading bacteria (11). Under certain circumstances p62 can also be degraded by endosomal microautophagy (12), but it together with ubiquitinated cargos is primarily degraded during selective autophagy through the interaction with FIP200 and the subsequent mutually exclusive interaction with LC3 (13–15). p62 is also known as a multifunctional signaling hub, as it participates in the activation of mechanistic target of rapamycin complex 1 (mTORC1) in nutrient sensing, NF-κB activation during inflammation, apoptosis and the activation of the Keap1-Nrf2 pathway for antioxidant response, as well as its mentioned adaptor role in selective autophagy (16). Among them, the p62-mediated Nrf2-pathway is coupled with selective autophagy specifically. Nuclear factor erythroid 2–related factor 2 (Nrf2) is a basic leucine zipper (bZIP) transcription factor, and its heterodimer with small MAF proteins controls the expression of proteins that protect against oxidative damage triggered by injury and inflammation. The tumor suppressor Kelch-like ECH-associated protein 1 (Keap1) is an adaptor of cullin3-based ubiquitin ligase for Nrf2. When p62 is phosphorylated at Ser349 under selective autophagy conditions, the S349-phosphorylated p62 interacts with Keap1 in competition with Nrf2, resulting in the inactivation of Keap1 and subsequent Nrf2-activation.

Recent growing lines of evidence shed light on unique feature of p62, phase-separation of p62 dependent on the presence of ubiquitin chains (17). The phase-separated p62 droplets allow the exchange of their components, including ubiquitin, LC3 and Keap1, with the surrounding environment (17–20). Inside a liquid-like droplet, molecules would be predicted to maintain their conformation and activity. Consequently, the droplets could also serve as nodes, from which signaling cascades could be activated in the context of selective autophagy and the Keap1-Nrf2 pathway (20,21). p62 clustering appears to require multiple ubiquitin chains of at least three ubiquitin moieties. Various ubiquitin chain linkages, but especially K63, are able to promote clustering, whereas free monoubiquitin or unanchored ubiquitin chains, specifically the K48 linkage, inhibit p62 clustering (18). These p62 structures arise from existing p62 filaments that are cross-linked by polyubiquitinated substrates (21,22). Inside these clustered droplets, p62 appears to retain little mobility, whereas ubiquitin, LC3 and Keap1 may be able to diffuse more easily within the cluster and the surrounding cytosol (17,18,20). Finally, cluster formation is rendered more efficient by the presence of NBR1, another autophagic adaptor that cooperates with p62 in cells (18,19). Thus, p62-droplets and/or gels are not simple substrates for autophagy but serve as platforms for both autophagosome formation and anti-oxidative stress (20,21). Whereas some ALS/FTD-related LIR or KIR p62 mutants showed the decreased Nrf2-activation or the delayed autophagic degradation (Table 1), there is no consensus view how mutations with different functional defects cause ALS/FTD. Herein, we investigated the effects of ALS/FTD-related p62 mutations in LIR and KIR on the Keap1-Nrf2 pathway, autophagic turnover, and the droplet dynamics and showed that a common defect in all p62 mutants is slower inner fluidity of the droplets.

**Table 1.** Functional defects of p62-mutants on the basis of previous reports.

| Mutation of p62<br>(Related-diseases) | Keap1-interaction<br>(Nrf2-activation) | LC3-interaction (autophagic<br>degradation of p62) | Ref. |
| --- | --- | --- | --- |
| D336N<br>(ALS/FTD, PDB) | n.d. (n.d.) | n.d. (n.d.) | – |
| D337E<br>(ALS/FTD) | n.d. (n.d.) | + (n.d.) | (25,26) |
| L341V<br>(ALS) | + (+) | – (–) | (26,27) |
| K344E<br>(ALS/FTD) | + (+) | n.d. (n.d.) | (27) |
| P348L<br>(ALS/FTD) | – (–) | n.d. (n.d.) | (27) |
| S349T<br>(PDB) | +/- (n.d.) | n.d. (n.d.) | (28) |
| G351A<br>(ALS/FTD) | – (–) | n.d. (n.d.) | (27) |

n.d.: not determined., +/-: less interaction comparable to wild-type p62, +: interaction or degradation comparable to wild-type p62. c.1006G>A (p.Asp336Asn) was found in ClinVar (https://www.ncbi.nlm.nih.gov/clinvar/variation/542160/).

## Results

### Binding models of disease-related p62 mutants with LC3 and Keap1

Disease-related p62 mutants, which have missense mutations locating at LIR to KIR (Fig. 1A) have been investigated their effects on selective autophagy and the p62-mediated Nrf2 activation (Table 1), but the consistent effect among these mutations was not specified. We previously solved the binding modes of the LIR with LC3 (PDB 2ZJD) and of the KIR with Keap1 (PDB 3ADE). On the basis of these structural data, we constructed models of the p62 mutants-LC3 or −Keap1 complex (Fig. 1B-D). As shown in Figure 1B, p62 Asp336 forms electrostatic interaction with LC3 Arg11. Loss of negative charge by D336N mutation will weaken this interaction and thereby weaken the interaction with LC3. On the other hand, D337E mutation will strengthen the interaction with LC3 by creating electrostatic interaction between Glu337 and Lys49 (Fig. 1B) (Asp337 is located distally from Lys49 and thus cannot form electrostatic interaction). The side-chain of Leu341 is bound to the L-site of LC3, which will be weakened by L341V mutation by worse fitting (Fig. 1C). P348L and G351A mutation will weaken the interaction with Keap1 by causing steric clash with Tyr525 and Tyr572 of Keap1, respectively (Fig. 1D). On the other hand, T350A mutation may weaken the interaction with Keap1 indirectly as following mechanism: The side-chain of Thr350 forms an intramolecular hydrogen-bond with the main-chain of Glu352 (Fig. 1D). T350A mutation impairs this hydrogen-bond, which may cause a conformational change in p62 KIR, leading to the reduced affinity with Keap1.

**Figure 1.**
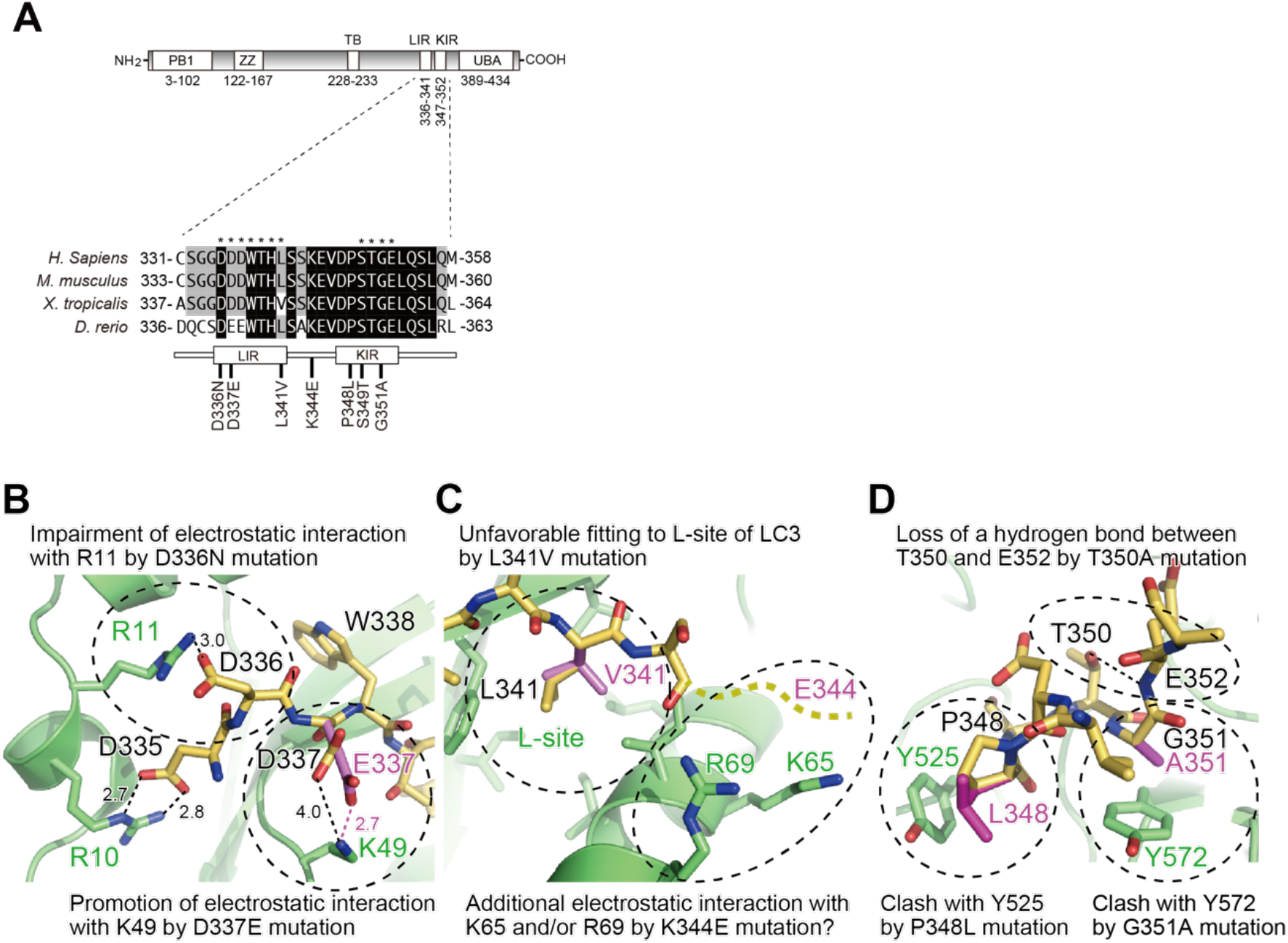
Binding models of disease-related p62 mutants with LC3 and Keap1. **(A)** Domain structure of p62 and the disease-related mutations in LIR and KIR. PB1: Phox and Bem1p. ZZ: Zinc finger. TB: TRAF6-binding domain. LIR: LC3-interacting region. KIR: Keap1-interacting region. UBA: Ubiquitin-associated. **(B)** Effect of D336N and D337E mutation on the interaction with LC3. The side-chain of Glu337 was manually modeled. Numbers indicate distance in Å. **(C)** Effect of L341V and K344E mutation on the interaction with LC3. The side-chain of Val341 was manually modeled. Figures in B and C were prepared using the crystal structure of the LC3B-p62 LIR complex (PDB 2ZJD). Residue numbers refer to human proteins. **(D)** Effect of P348L, T350A, and G351A mutation on the interaction with Keap1. Figure in D was prepared using the crystal structure of the Keap1-p62 KIR complex (PDB 3ADE). Residue numbers refer to human proteins.

### Interaction of disease-related p62 mutants with FIP200, LC3 and Keap1

To test whether the disease-related p62 mutants have an effect on selective autophagy and the Keap1-Nrf2 system, we generated *p62*-deficient HEK293T cells (Supplementary Fig. S1) and expressed each FLAG-tagged wild-type and p62 mutant. Subsequently, the cell lysates were conducted to SDS-PAGE, followed by immunoblot analysis. Remarkably, while we did not observe any significant difference in total level of p62 in the cells expressing mutants, some p62 mutants, in particular p62^K344E^ mutant underwent higher phosphorylation at S349 than wild-type p62 (Fig. 2A). The levels of Keap1, LC3-II and FIP200 were not influenced by the expression of p62 mutant proteins (Fig. 2A). Since the phosphorylation at S349 of p62 enhances the binding affinity to both Keap1 and FIP200 (15,23), the higher binding affinity to FIP200 and Keap1 of p62^K344E^ mutant was expected. But, immunoprecipitation assay with anti-FLAG antibody showed that mutants except a disease-related mutant p62^G351A^ and a Keap1-binding defective p62^T350A^ (24) have a binding affinity to Keap1 comparable to wild-type p62 (Fig. 2B). The immunoprecipitates prepared in our experimental settings did not contain FIP200 (data not shown). Consistent with the structural models (Fig. 1B and C), p62^D337E^ and p62^K344E^ mutants showed higher affinity to both LC3-I and II forms (Fig. 2B). The affinity of p62^P348L^ to LC3 was also higher (Fig. 2B). Meanwhile, p62^D336N^, p62^L341V^ and LIR mutant p62^W338A^ ^L341A^ (14) revealed decreased affinity (Fig. 2B). These results on LC3- and Keap1-interactions with p62 were consistent with those reported by other groups (25–28) (Tables 1 and 2).

**Table 2.** Summary of interaction of p62 mutants with Keap1 and LC3.

| Mutation of p62<br>(Related-diseases) | Keap1-interaction<br>(IP and NanoBRET) | LC3-interaction<br>(IP and NanoBRET) | FIP200-interaction<br>(NanoBRET) |
| --- | --- | --- | --- |
| D336N<br>(ALS/FTD, PDB) | + and + | – and + | + |
| D337E<br>(ALS/FTD) | + and + | ++ and + | + |
| L341V<br>(ALS) | + and – | – and – | – |
| K344E<br>(ALS/FTD) | + and + | ++ and + | + |
| P348L<br>(ALS/FTD) | + and – | + and + | + |
| S349T<br>(PDB) | + and + | + and + | + |
| G351A<br>(ALS/FTD) | – and – | + and + | + |
| W338A L341A<br>(–) | + and + | – and – | + |
| T350A<br>(–) | – and – | + and + | + |

–: weak or not detectable, +: interaction comparable to wild-type p62, ++: higher affinity than wild-type p62.

**Figure 2.**
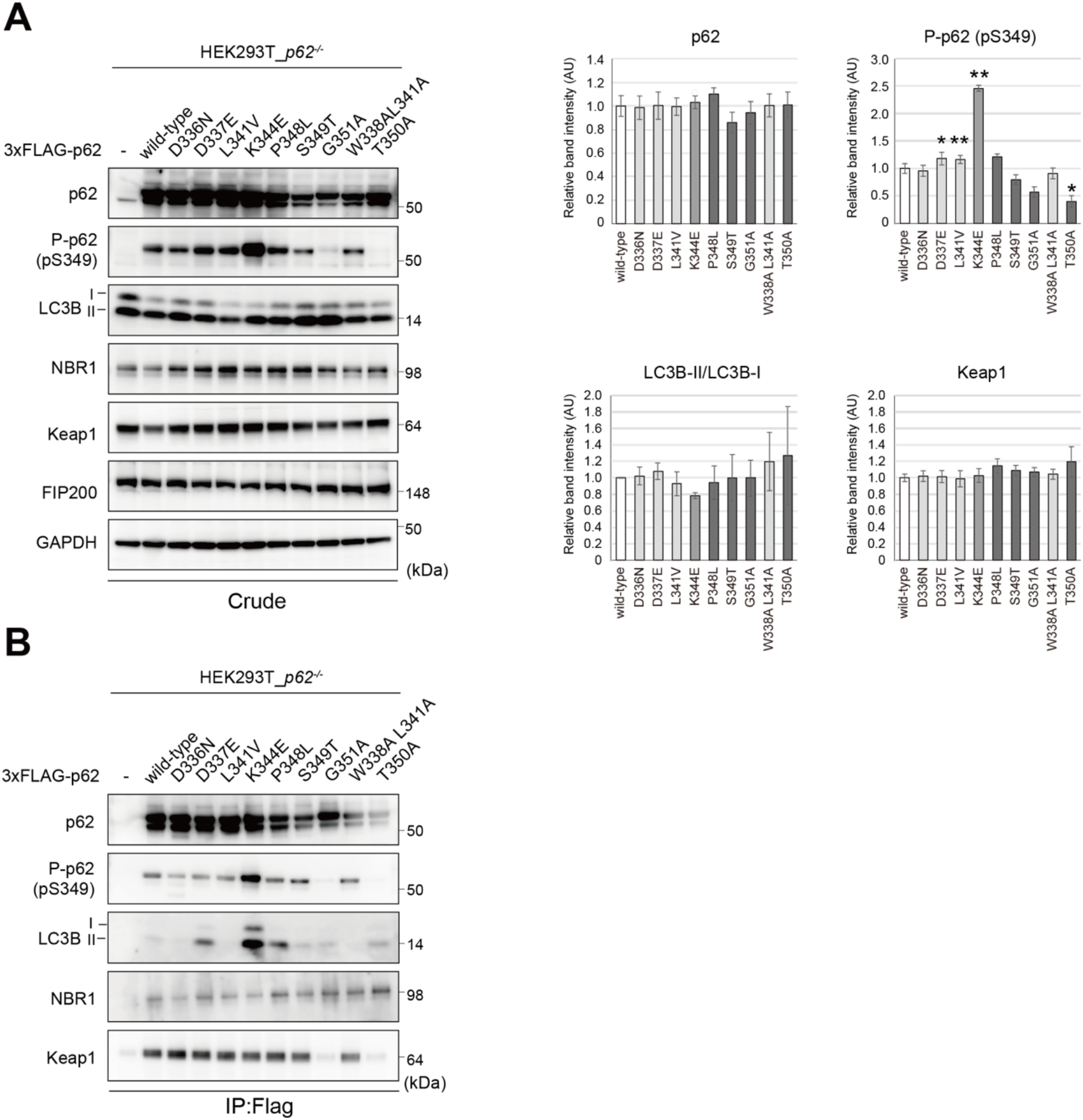
Interaction of disease-related p62 mutants with the binding partners. **(A)** Immunoblot analysis. FLAG-tagged wild-type and mutant p62 were expressed in *p62^−/−^* HEK293T cells. At 24 hr after transfection, cell lysates were prepared and subjected to immunoblot analysis with the indicated antibodies. Data shown are representative of four separate experiments. Bar graphs indicate the quantitative densitometric analysis of the indicated proteins relative to GAPDH (n = 4). Data are means ± s.e. \**p* < 0.05, and \*\**p* < 0.01 as determined by two-sided Welch’s *t*-test. **(B)** Immunoprecipitation assay. Cell lysates prepared as described in (A) were immunoprecipitated with FLAG antibody, and the immunoprecipitants were subjected to immunoblot analysis with the indicated antibodies. Data shown are representative of three separate experiments.

### In vivo interaction of disease-related p62 mutants with FIP200, LC3 and Keap1

The p62 is present as gels in cells (20), raising a possibility that the results based on the immunoprecipitation assay do not reflect *in vivo* dynamics. To investigate the effect of p62 mutations on their affinity to the binding partners in cells, we conducted a bioluminescence resonance energy transfer (BRET)-based assay that uses NanoLuc® Luciferase as the BRET energy donor and HaloTag® protein labeled with the HaloTag® NanoBRET™ 618 fluorescent Ligand as the energy acceptor to measure the interaction of two binding partners in live cells (29) (Fig. 3A). LC3B, one of three LC3 homologue and FIP200 were fused with NanoLuc Luciferase at the N-terminus, and Keap1 were fused it at the C-terminus. p62 was fused with HaloTag at the N-terminus. These constructs were transfected into *p62*-deficient HEK293T cells, and we confirmed efficient expression of each protein by immunoblot analysis (Fig. 3B). NanoBRET assay revealed interaction features of p62 distinct to the immunoprecipitation assay (Fig. 3C-E). While the immunoprecipitation assay revealed that disease-related mutants except for p62^G351A^ interact with Keap1 to the same extent (Fig. 2B), the NanoBRET assay showed that in addition to p62^G351A^, p62^L341V^ and p62^P348L^ also have less ability to bind to Keap1 (Fig. 3C), which is agreement with the structural models (Fig. 1D). In the case of the interaction of p62 with LC3B, the affinity of p62^L341V^ and p62^W338A^ ^L341A^ to LC3B was less compared with that of wild-type p62 (Fig. 3D). In contrast to the immunoprecipitation assay, no p62 mutants showed increased binding avidity to LC3B in NanoBRET (Fig. 3D). Though we did not detect any interaction of p62 with FIP200 in the immunoprecipitation assay (data not shown), the interaction was detectable in the NanoBRET assay (Fig. 3E). p62^L341V^ mutant exhibited slightly but significant defect in the interaction with FIP200 (Fig. 3E).

**Figure 3.**
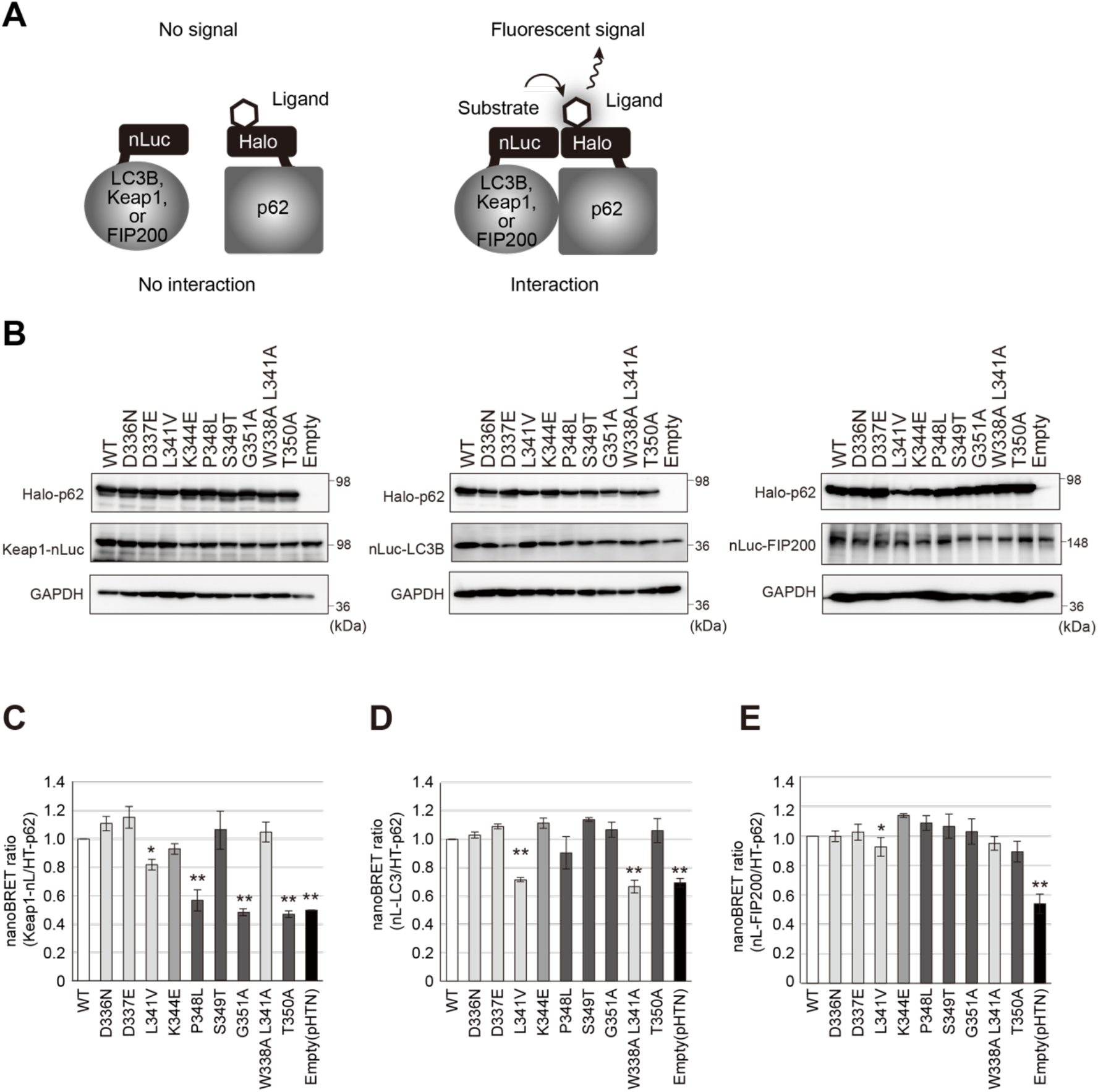
*In vivo* interaction of disease-related p62 mutants with the binding partners. **(A)** Principal of NanoBRET assay. **(B)** Immunoblot analysis. Each HaloTag-tagged wild-type and mutant p62 was co-expressed with NanoLuc Luciferase-tagged Keap1, LC3 or FIP200 in *p62^−/−^* HEK293T cells. At 24 hr after the transfection, cell lysates were prepared and subjected to immunoblot analysis with the indicated antibodies. Data shown are representative of four separate experiments. (**C**-**E**) NanoBRET assay between p62 and Keap1 (C), LC3B (D) and FIP200 (E). Bar graphs indicate the quantitative analysis of the indicated mutant p62 proteins relative to wild-type p62 (n = 3). Data are means ± s.e. \**p* < 0.05, and \*\**p* < 0.01 as determined by two-sided Welch’s *t*-test.

### The effects of p62 mutants on cellular functions

p62 serves as a positive regulator of Nrf2 through the direct interaction with Keap1(23,24). The stress response through Nrf2 is universal among tissues, but it plays pivotal roles in livers where are responsible for a production in phase-I, phase-II drug-metabolizing enzymes and efflux transporters (30). To evaluate the effect of each mutation on the p62-mediated Nrf2-activation at high sensitivity, we developed adenovirus-vectors, which are able to express target proteins effectively even in primary cultured cells, for wild-type human p62 and each mutant. We infected them with primary mouse culture hepatocytes and confirmed their effective expressions (Fig. 4A). In comparison with lacZ-expressing primary hepatocytes, the amount of nuclear Nrf2 protein increased upon the expression of wild-type p62 (Fig. 4A). Such effect was hardly observed in the case of p62^P348L^, p62^G351A^ and p62^T350A^ (Fig. 4A). In good agreement with these results, real-time PCR analysis revealed that while gene expression of Nrf2-targets such as *Glutathione S-transferase Mu 1* (*Gstm1*), *UDP-glucose 6-dehydrogenase* (*Ugdh*) and *NAD(P)H dehydrogenase, quinone 1* (*Nqo1*) was markedly induced by overexpression of wild-type p62, but not the G351A, P348L and T350A mutants (Fig. 4B). We did not find any defect in the level of nuclear Nrf2 and the induction of Nrf2-target genes upon expression other disease-related mutants (Fig. 4A and B).

**Figure 4.**
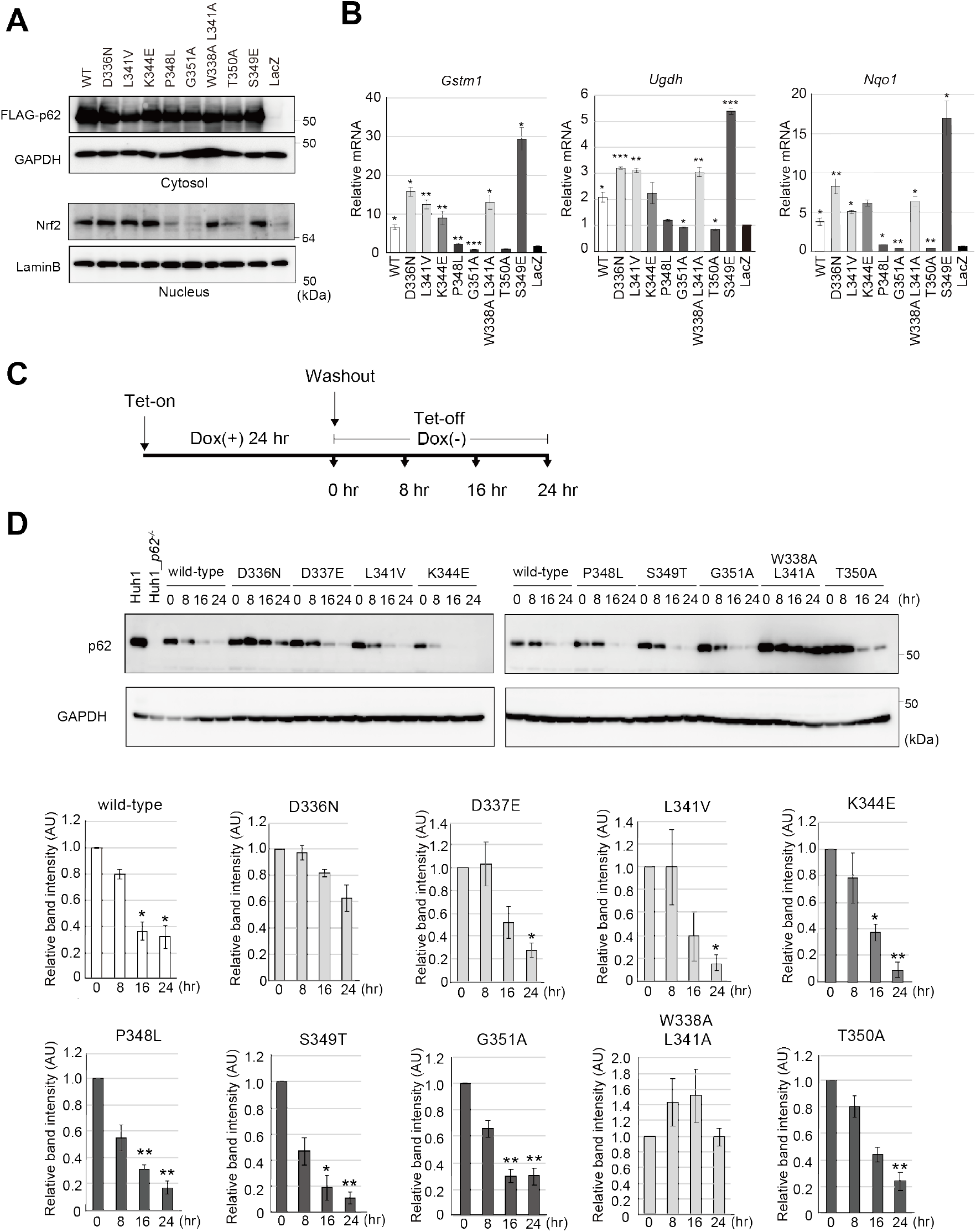
Nrf2-activation by and the degradation of disease-related p62. **(A)** Immunoblot analysis. Each wild-type p62 and the mutants was expressed in primary mouse hepatocytes using adenovirus system. 48 hr after the infection, the cells were fractionated into cytosolic and nuclear fractions, which were subjected to immunoblot analysis with the indicated antibodies. Data shown are representative of three separate experiments. **(B)** Gene expression of Nrf2-targets. Total RNAs were prepared from primary mouse hepatocytes expressing LacZ (n` = 3), wild-type p62 (n = 3), each mutant (n = 3). Values were normalized against the amount of mRNA in LacZ-expressing hepatocytes. RT qPCR analyses were performed as technical replicate on each biological sample. Data are means ± s.e. \**p* < 0.05, \*\**p* < 0.01, and \*\*\**p* < 0.001 as determined by two-sided Welch’s *t*-test. **(C)** Schematic representation of experiments conducted to monitor p62 protein level. **(D)** Immunoblot analysis. *p62*-knockout Huh-1 cells were expressed FLAG-p62 or each disease-related mutant by the treatment of doxycycline (DOX) for 24 hr and then cultured with media in absence of DOX for indicated time. The cell lysates were prepared, and then subjected to SDS-PAGE followed by immunoblotting with the indicated antibodies. Data are representative of three separate experiments. Bar graphs indicate the quantitative densitometric analysis of the indicated proteins relative to whole proteins estimated by GAPDH (n = 3). Data are means ± s.e. \**p* < 0.05, and \*\**p* < 0.01 as determined by two-sided Welch’s *t*-test.

In the next series experiments, we investigated degradation of each disease-related p62 mutant protein. To do this, we developed *p62*-knockout Huh-1 cells (Supplementary Fig. S1) and introduced with two regulator gene cassettes, CAG-rtTA and TRE-FLAG-fused wild type and a series of mutant p62. To monitor the half-live of wild-type and each mutant p62, FLAG-tagged wild-type or each mutant was expressed by the treatment of DOX for 24 hr, and then the cells were cultured in medium without DOX for indicated time (Fig. 4C). While the amount of wild-type FLAG-p62 decreased by approximately 60% at 24 hr after removal of DOX, that of p62^D336N^ and of p62^W338A^ ^L341A^ remained unchanged (Fig. 4D). Unexpectedly, L341V mutant, which exhibited the loss of binding ability to LC3 in both the immunoprecipitation and NanoBRET assays (Fig. 2B and Fig. 3D) was degraded to same extent with wild-type p62 (Fig. 4D). Likewise, similar to wild-type p62, the levels of other disease-related mutants including p62^D337E^, p62^K344E^, p62^P348L^, p62^S349T^ and p62^G351A^ were significantly declined after removal of DOX (Fig. 4D).

### Decreased inner-fluidity of disease-related p62-liquid droplets

Mutants of p62, which cause same disease(s), showed separate defects in cell physiology (Fig. 4). What is a common feature of disease-related p62 mutant proteins? We hypothesized that the mutations of p62 in LIR and KIR affect the formation and/or dynamics of p62 liquid-droplets. Each GFP-tagged wild-type and mutant p62 was expressed in *p62*-deficient Huh-1 cells by the treatment of DOX for 48 hr, and we observed the GFP-p62 signal. A number of GFP-p62- positive structures regardless of the presence or absence of mutation were observed. We first measured the circularity of the p62-positive structures. The closer circularity is to 1, it implies an ability to form a liquid droplet. As shown in Figure 5A, in comparison with the circularity of wild-type p62-structures, that of most mutant p62-structures had closer value to 1. Even in the case of structures with diameter over 1.5 μm, we did not observe any less circularity of mutant p62-structures (Fig. 5A), suggesting that the disease-related mutants become liquid droplets. Time-lapse imaging showed that the p62-structures with or without mutations moved through the cytoplasm and occasionally fused with each other (Supplementary Movies S1-S10), matching with criteria of liquid droplets. We also evaluated the number and size in p62-liquid droplets and summarized in Table 3. The number and size of droplets consisting of p62^L341V^ were larger than and comparable to those of wild-type droplets (Fig. 5B and C). While the number of p62^K344E^ droplets was comparable to that of wild-type droplets, the size was larger (Fig. 5B and C). These results indicate that both mutants are prone to form liquid droplets. By contrast, the number and size of p62^S349T^- and p62^G351A^-droplets were less than those of wild-type droplets (Fig. 5B and C), suggesting less ability to form liquid droplets. Finally, to investigate the inner fluidity of p62-droplets, we measured fluorescence recovery after photobleaching (FRAP) of wild-type and each mutant p62-droplet. The droplets with diameter with 1.5-2 μm were measured. Consistent with previous reports by us and other groups (17,18,20), the average fluorescence recovery time after photobleaching of GFP-p62-droplets is not so fast, 4.58 ± 0.37 min (Fig. 5D, Supplementary Figure S2 and Supplementary Movie S11). Remarkably, all droplets formed by disease-related p62 mutants including PDB-associated one (S349T) as well as core LIR and KIR mutants displayed slower fluorescence recovery time in greater or lesser degrees (Fig. 5D, Supplementary Figure S2 and Supplementary Movie S11- 20). The mutant p62 droplets were divided into three groups; much slower (p62^D336N^ and p62^T350A^), mild slower (p62^D337E^, p62^L341V^, p62^K344E^, p62^G351A^ and p62^W338A^ ^L341A^) and a few slower (p62^P348L^ and p62^S349T^) (Table 3). The magnitude of each p62-droplet in the slower fluidity was not correlated to the localization of LC3 and/or Keap1 to the droplets (Supplementary Fig. S3A, B and Table 3).

**Table 3.** Summary of circularity, number and size, and inner fluidity of p62-liquid droplets.

| Mutation of p62<br>(Related-diseases) | Circularity<br>(1-2 $\mu\text{m}$ and $>1.5\mu\text{m}$) | Number and size | Inner fluidity |
| --- | --- | --- | --- |
| D336N<br>(ALS/FTD, PDB) | ++ and ++ | ++ and – | --- |
| D337E<br>(ALS/FTD) | ++ and + | ++ and – | -- |
| L341V<br>(ALS) | ++ and ++ | ++ and + | -- |
| K344E<br>(ALS/FTD) | + and + | + and ++ | -- |
| P348L<br>(ALS/FTD) | ++ and ++ | ++ and – | – |
| S349T<br>(PDB) | ++ and ++ | – and – | – |
| G351A<br>(ALS/FTD) | + and + | – and – | -- |
| W338A L341A<br>(–) | ++ and + | ++ and – | -- |
| T350A<br>(–) | + and + | ++ and – | --- |

–: fewer or lower, +: comparable to wild-type p62, ++: larger. D336N and W338L L341V mutants show delayed autophagic degradation.

**Figure 5.**
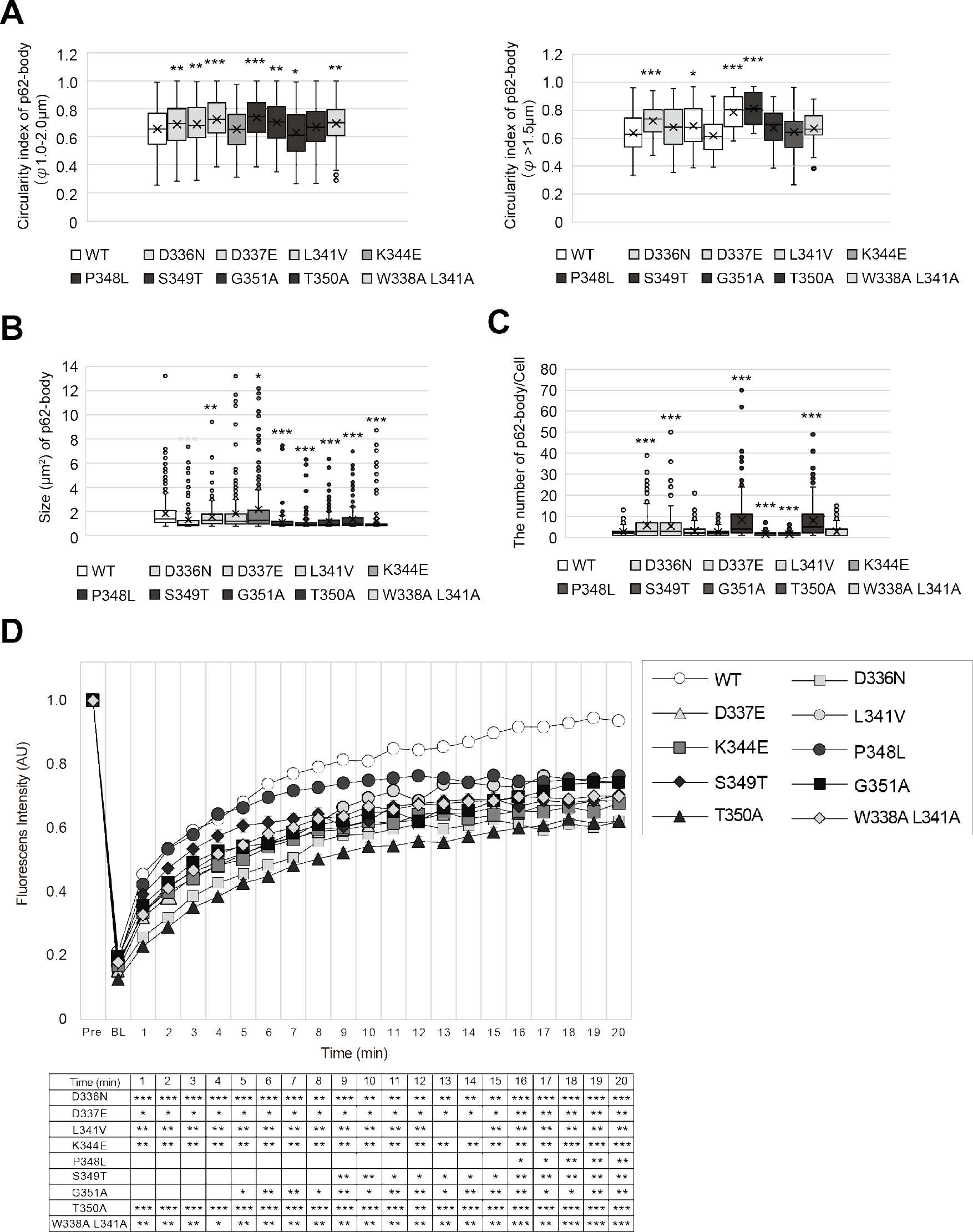
Droplets of disease-related p62. (**A-C**) Circularity (A), number (B) and size (C) of p62-structures. *p62*-knockout Huh-1 cells were expressed GFP-p62 or each disease-related mutant by the treatment of doxycycline (DOX) for 24 hr. The circularity, number and size of GFP-p62-structures were determined by procedures described in Experimental procedures. (**D**) FRAP. Huh-1 cells were transfected by GFP-p62 and the mutants. 24 hr after the transfection, the p62-bodies labelled by GFP-p62 liquid droplets were photobleached, and then time of fluorescent recovery was measured. Data are means ± s.e of non-photobleached (n = 10) and photobleached (n = 10) dots. \*\**p* < 0.01, and \*\*\**p* < 0.001 as determined by two-sided Welch’s *t*-test. Bar: 2 μm.

## Discussion

Past a few years, a growing body of evidence indicated the formation and subsequent gel or aggregate transition of liquid droplets consisting of distinct ALS/FTD-related mutant proteins including FUS (31–33), TDP-43 (34), hnRNPA1 (35) and Tau (36) as well as dipeptide repeat derived from C9orf72 gene (37,38). On the other hand, whether disease-related p62 mutants exhibit similar abnormality or not remained unclear. All ALS/FTD-associated p62 mutant proteins, which harbor missense mutations in LIR and KIR exhibited increased or decreased binding affinity to their binding partners, LC3 and Keap1 (Figs. 2 and 3), and some of them but not all resulted in suppression of the p62-mediated Nrf2-activation and of the autophagic turnover (Fig. 4). However, such defects were in cases of some mutations, and not consistent among all mutations. Meanwhile, while their ability to form droplets, a consistent abnormality among the mutant proteins was slower inner fluidity of the droplets (Fig. 5), probably due to deviation from proper multivalency. Thus, we propose that a primary defect in ALS/FTD with p62 missense mutations is not due to defect at least in selective autophagy and anti-oxidative stress response, rather it is attributed to a qualitative change of p62-liquid droplets; slower inner fluidity of the droplets followed by the aggregation. Though we observed functional defects of only some restricted p62 mutant proteins in the activation of Nrf2 and autophagic turnover at least in short period, prolonged qualitative change of the p62-droplets should be accompanied by those functional defects and/or aggregation-dependent gain-of-function cytotoxic effects. Actually, the aggregated structures such as aggregate-prone Ape1-complex is not recognized as a selective substrate due to a failure of an adaptor protein to function on the aggregates (39). Keap1 traffics from cytoplasm to the droplets (20) and is sequestered into disease-associated p62-aggregates (20,40,41). Our hypothesis is also supported by a fact that missense mutations of *SQSTM1/p62* related to ALS/FTD have been identified throughout the gene including regions encoding the intrinsically disordered region and UBA domain, both of which are dispensable for the liquid-droplet formation.

### Experimental procedures

#### Cell culture

Huh-1 (JCRB0199) and HEK293T (ATCC CRL-3216) cells were grown in Dulbecco’s modified Eagle medium (DMEM) containing 10% fetal bovine serum (FBS), 5 U/ml penicillin, and 50 μg/ml streptomycin. For overexpression experiments, Huh-1 and HEK293T cells were transfected with Lipofectamine 3000 (Thermo Fisher Scientific, Waltham, MA, USA) and Polyethylenimine 25kD linear (PEI, #23966, Polysciences), respectively. To generate *p62*- knockout HEK293T cells, *p62* guide RNA designed using the CRISPR Design tool (http://crispr.mit.edu/) was subcloned into pX330-U6-Chimeric_BB-CBh-hSpCas9 (Addgene #42230), a human codon-optimized SpCas9 and chimeric guide RNA expression plasmid. The HEK293T cells were transfected with vectors pX330 and cultured for 2 days. Thereafter, the cells were sorted and expanded. Loss of *p62* was confirmed by immunoblot analysis with anti-p62 antibody. All cells were authenticated by STR profile and tested for mycoplasma contamination.

### Isolation of mouse primary hepatocytes

Eight-week-old mice were anesthetized with isoflurane, and the mouse livers were perfused by reverse flow through the vena cava using 20 mL of ex-perfusate solution (pH 7.4, 140 mM NaCl, 5mM KCl, 0.5 mM NaH2PO4, 10 mM HEPES, 4mM NaHCO_3_, 50 mM glucose and 0.5 mM GEDTA) and 30 mL of perfusate solution (pH 7.6, 70 mM NaCl, 6.7 mM KCl, 100 mM HEPES, 5 mM CaCl2, 35 units collagenase). After dissection, hepatocytes suspension was washed three times in HBSS (Gibco, Thermo Fisher Scientific) and the cells were grown in collagen-coated plates with William’s Medium E (Gibco, Thermo Fisher Scientific) supplemented with 10% fetal bovine serum (FBS).

### Adenovirus infection

Wild-type p62 and mutant adenoviruses were prepared using the Adenovirus Expression Vector Kit (TAKARA BIO., Otsu, Japan). To express exogenous p62 and the mutant p62 proteins in mouse primary culture hepatocytes, the cells were plated onto 6-well dishes. At 24 hr after plating, the medium was replaced with fresh medium containing adenovirus at a multiplicity of infection (MOI) of 100.

### Generation of Tet-ON cells

Tetracycline-mediated p62-expressing cell lines were generated using the reverse tet-regulated retroviral vector. A cassette consisting of the gene encoding packaging signal (ψ), the reverse tetracycline controlled transactivator (rtTA), the internal ribosome entry site from the ribosome (IRES), the blasticidin S deaminase (BSD), a heptamerized tet operator sequence (tetO), the minimal human cytomegalovirus immediate early promoter designated Phcmv*-1 (CMV), and human p62 cDNA or various p62 mutants were cloned into a Moloney murine leukemia virus (M-MuLV) retroviral vector pLXSN back bone. Retrovirus packaging cells, PLAT-E, transfected with the vectors were cultured at 37°C for 24 hr. After changing the medium, the virus producing PLAT-E was further incubated at 37°C for 24 hr. The viral supernatant was collected and used immediately for infection. p62-deficient Huh-1 were plated on 35 mm dishes in 3 mL growth medium at 24 hr before infection. Just before infection, the medium was replaced with undiluted viral supernatant with 8 μg/ml polybrene (Sigma). 24 hr later, the cells were introduced into the selection medium containing 2 μg/ml of Puromycin Dihydrochloride (Fujifilm Wako Pure Chemical Corporation). The cells remaining after 5 days were used in the experiments. To induce the expression of p62, the cells were treated with 250 ng/ml of doxycycline (Dox, Sigma) for 24 hr.

### Immunoblot analysis

Cells were lysed in ice-cold TNE buffer (50 mM Tris-HCl, pH 7.5, 150 mM NaCl, 1 mM EDTA) containing 1% Triton X-100 and protease inhibitors. Nuclear and cytoplasmic fractions were prepared using the NE-PER Nuclear and Cytoplasmic Extraction Reagents (Thermo Fisher Scientific). Samples were separated by SDS-PAGE and then transferred to polyvinylidene difluoride (PVDF) membranes. For immunoprecipitation analysis, cells were lysed by 200 μl of TNE, and the lysate was then centrifuged at 10,000 x *g* for 10 min at 4°C to remove debris. Then, 800 μl of TNE and 1 μg of anti-FLAG antibody (M185-3L, Medical & Biological Laboratories) were added to the lysate, and the mixture was mixed under constant rotation for 12 hr at 4°C. The immunoprecipitates were washed five times with ice-cold TNE. The complex was boiled for 10 min in SDS sample buffer in the presence of 2-mercaptoethanol to elute proteins and centrifuged at 10,000 x *g* for 5 min. Anti-S349-phosphorylated p62 polyclonal antibody was raised in rabbits by using the peptide Cys+KEVDP(pS)TGELQSL as an antigen (23). Antibodies against p62 (PM066, Medical & Biological Laboratories), LC3B (#2775, Cell Signaling Technology), GAPDH (6C5, Santa Cruz Biotechnology), FIP200 (17250-1-AP, Proteintech Group), Lamin B (M-20, Santa Cruz Biotechnology), Nrf2 (H-300; Santa Cruz Biotechnology) and Keap1 (10503-2-AP; Proteintech Group) were purchased from the indicated suppliers and employed at 1:500 dilution. Blots were then incubated with horseradish peroxidase-conjugated secondary antibody (Goat Anti-Mouse IgG (H + L), 115-035-166, Jackson ImmunoResearch; Goat Anti-Rabbit IgG (H + L) 111-035-144; Goat Anti-Guinea Pig IgG (H + L)) and visualized by chemiluminescence.

### NanoBRET assay

HEK293T *p62*-knockout cells were co-transfected with Halo-tagged p62 and NanoLuc-tagged Keap1, LC3B or FIP200. 24 hr after the transfection, the cells were seeded into a white 96-well plate (152028, Thermo Fisher Scientific) at 2 x10^4^ cells/well and labeled with Halo tag NanoBRET 618 ligand (G9801, Promega) for 24 hr. Thereafter, the cells were incubated with 50 μl of DMEM containing 4% FBS and NanoBRET substrate (N1571, Promega) for 5 min. The BRET signal was measured by a GloMax® Discover Microplate Reader (GM3000, Promega) using a 460-nm filter for the donor signal and a 610-nm filter for the acceptor signal. The BRET ratio was determined by dividing the acceptor signal by the donor signal.

### Immunofluorescence microscopy

Cells grown on coverslips were fixed in 4% paraformaldehyde in PBS for 10 min, permeabilized with 0.1% Triton X-100 in PBS for 5 min, blocked with 0.1% (w/v) gelatin (Sigma-Aldrich) in PBS for 45 min, and then incubated overnight with primary antibodies diluted 1:200 in gelatin/PBS. After washing, cells were incubated with Goat anti-Guinea pig IgG (H + L) Cross-Adsorbed Secondary Antibody, Alexa Fluor 488 (A11073, Thermo Fisher Scientific), and Goat anti-Mouse IgG (H + L) Highly Cross-Adsorbed Secondary Antibody, Alexa Fluor 647 (A21236, Thermo Fisher Scientific) at a dilution ratio of 1:1000 for 60 min. Cells were imaged using a confocal laser-scanning microscope (Olympus, FV1000) with a UPlanSApo ×60 NA 1.40 oil objective lens.

### Determination of the number, size and circularity of p62-bodies

GFP-p62 were expressed into *p62* knockout Huh-1 cells using dox-inducible system. The cells were fixed with 4% paraformaldehyde in PBS for 10 min, followed by Hoechst staining. Cells were imaged using Confocal Quantitative Image Cytometer (CQ1, Yokogawa). To quantify the number, size, and circularity of p62-droplets, the images over 300 cells with GFP-signal were analyzed by Cellpathfinder software (Ver.R3.04, Yokogawa). To recognize p62-droplets exclusively, the GFP-positive structures with 0.8-5.0 μm in diameter were analyzed.

### Fluorescence recovery after photobleaching

To measure the fluorescence recovery after photobleaching (FRAP), cells were grown in 35mm glass base dishes (Iwaki, Japan). p62-bodies were bleached for 3 sec using a laser intensity of 70% at 480 nm, and then the recovery was recorded for the indicated time. After image acquisition, contrast and brightness were adjusted using Photoshop 2021v25.0 (Adobe).

### Quantitative real-time PCR (qRT-PCR)

cDNAs were synthesized with 1 μg of total RNA using FastGene Scriptase Basic cDNA Synthesis (NE-LS62, NIPPON Genetics). Quantitative PCR was performed with TaqMan® Fast Advanced Master Mix (444556, Thermo Fisher Scientific) on a QuantStudio^TM^ 6 Pro (A43180, Thermo Fisher Scientific). Signals were normalized against *Gusb* (β-glucuronidase). Predesigned TaqMan Gene Expression Assays including primer set and TaqMan probe (Gusb; Mm01197698_m1, Nqo1; Mm01253561_m1, Ugdh; Mm00447643_m1 and Gstm1; Mm00833915_g1 were purchased from Thermo Fisher Scientific.

## Data availability

The authors declare that the data supporting the findings of this study are available within the article and its supporting information.

## Supporting information

This article contains supporting information.

## Author Contributions

Masaaki K. and Y.I. designed and directed the study. F.O., Y.I., S.K., AH.E-G., Y-s.S. and Masato K.carried out the biochemical and cell biological experiments. Masaaki K. wrote the manuscript. All authors discussed the results and commented on the manuscript.

## Funding and additional information

F.M.O is supported by Japanese Government (Monbukagakusho: MEXT) Scholarship. Y.I. is supported by Grant-in-Aid for Scientific Research (C) (20K06644). S.K. is supported by a Grant-in-Aid for Scientific Research (C) (20K06549). Y.-S.S. is supported by a Grant-in-Aid for Scientific Research (C) (19K15043). M.K.supported by Grant-in-Aid for Scientific Research on Innovative Areas (19H05706), Grant-in-Aid for Scientific Research (A) (21H004771), a Japan Society for the Promotion of Science (an A3 foresight program, to M.K.), and the Takeda Science Foundation (to M.K.). AH.E-G. would like to thank the Egyptian Government for financial support to carry out this research project.

